# Global Increases in Human Immunodeficiency Virus Neutralization Sensitivity Due to Alterations in the Membrane-Proximal External Region of the Envelope Glycoprotein Can be Minimized by Distant State 1-Stabilizing Changes

**DOI:** 10.1101/2021.11.01.466860

**Authors:** Qian Wang, Florian Esnault, Meiqing Zhao, Ta-Jung Chiu, Amos B. Smith, Hanh T. Nguyen, Joseph G. Sodroski

**Affiliations:** Department of Cancer Immunology and Virology, Dana-Farber Cancer Institute, Department of Microbiology, Harvard Medical School, Boston, MA 02215, USA; Department of Chemistry, University of Pennsylvania, Philadelphia, PA 19104, USA; Department of Immunology and Infectious Diseases, Harvard T. H. Chan School of Public Health, Boston, MA 02215, USA

**Keywords:** HIV-1 Env, MPER, native conformation, State 1, stabilization, cholesterol, membrane, triggerability

## Abstract

Binding to the receptor, CD4, drives the pretriggered, “closed” (State-1) conformation of the human immunodeficiency virus (HIV-1) envelope glycoprotein (Env) trimer ((gp120/gp41)_3_) into more “open” conformations. HIV-1 Env on the viral membrane is maintained in a State-1 conformation that resists binding and neutralization by commonly elicited antibodies. Premature triggering of Env before the virus engages a target cell typically leads to increased susceptibility to spontaneous inactivation or ligand-induced neutralization. Here, we show that single amino acid substitutions in the gp41 membrane-proximal external region (MPER) of a primary HIV-1 strain result in viral phenotypes indicative of premature triggering of Env to downstream conformations. Specifically, the MPER changes reduced viral infectivity and globally increased virus sensitivity to poorly neutralizing antibodies, soluble CD4, a CD4-mimetic compound, and exposure to cold. By contrast, the MPER mutants exhibited decreased sensitivity to the State-1 preferring inhibitor, BMS-806, and to the PGT151 broadly neutralizing antibody. Depletion of cholesterol from virus particles did not produce the same State 1-destabilizing phenotypes as MPER alterations. Notably, State 1-stabilizing changes in Env distant from the MPER could minimize the phenotypic effects of MPER alteration, but did not affect virus sensitivity to cholesterol depletion. Thus, membrane-proximal gp41 elements contribute to the maintenance of the pretriggered Env conformation. The conformationally disruptive effects of MPER changes can be minimized by distant State 1-stabilizing Env modifications, a strategy that may be useful in preserving the native pretriggered state of Env.

**IMPORTANCE:** The pretriggered shape of the human immunodeficiency virus (HIV-1) envelope glycoprotein (Env) is a major target for antibodies that can neutralize many strains of the virus. An effective HIV-1 vaccine may need to raise these types of antibodies, but this goal has proven difficult. One reason is that the pretriggered shape of Env is unstable and dependent on interactions near the viral membrane. Here we show that the membrane-proximal external region (MPER) of Env plays an important role in maintaining Env in a pretriggered shape. Alterations in the MPER resulted in global changes in Env conformation that disrupted its pretriggered shape. We also found that these disruptive effects of MPER changes can be minimized by distant Env modifications that stabilize the pretriggered shape. These modifications may be useful for preserving the native shape of Env for structural and vaccine studies.

## INTRODUCTION

The human immunodeficiency virus type 1 (HIV-1) envelope glycoprotein (Env) trimer mediates virus entry into host cells (1). HIV-1 Env is a Class I viral fusion protein composed of three gp120 exterior subunits and three gp41 transmembrane subunits (1–4). Env is synthesized in the rough endoplasmic reticulum where signal peptide cleavage, high-mannose glycan modification, and trimerization take place (5–7). This trimeric Env precursor (gp160) then traffics to the cell surface either through the conventional secretory pathway via the Golgi apparatus, where furin-mediated cleavage and modification with complex glycans occur, or through a pathway that bypasses the Golgi compartment (8–12). Envs transported to the cell surface through the Golgi apparatus are selectively incorporated into virions (8). On virions, Env samples at least three conformational states, reflecting its dynamic nature (13). Envs of primary HIV-1 strains mainly occupy a pretriggered, “closed” (State-1) conformation that resists the binding of most antibodies elicited during natural infection (13–17). More rarely elicited broadly neutralizing antibodies recognize conserved elements of the State-1 Env conformation (13–15, 18). Binding to the first receptor, CD4, triggers major conformational changes in Env, leading initially to a default intermediate conformation (State 2) and then to the full CD4-bound conformation (State 3) (19–25). The CD4-bound conformation (State 3) consists of a pre-hairpin intermediate in which the gp41 heptad repeat (HR1) region is exposed (26–28). Subsequent binding of the State-3 Env to the second receptor, either CCR5 or CXCR4, leads to the formation of an energetically stable gp41 six-helix bundle, a process that results in fusion of the viral and target cell membranes (29–42).

The gp41 transmembrane subunits anchor Env to the lipid membrane on the surface of infected cells and virus particles (43–45). The gp41 glycoprotein is composed of an N-terminal fusion peptide, a fusion peptide-proximal region (FPPR), two heptad repeat regions (HR1 and HR2), a membrane-proximal external region (MPER), a transmembrane region and a long cytoplasmic tail (43, 46). The gp41 MPER (residues∼659-683) plays different roles during the distinct stages of HIV-1 entry, which may be reflected in its dynamic structure. In current high-resolution structures of soluble or detergent-solubilized HIV-1 Env trimers, the MPER has been removed or is disordered (47–53). Lower-resolution tomograms of virion Envs have provided different views of the MPER conformation, from a stalk tightly organized near the trimer axis to a tripod with various degrees of splay (21, 54–57). Synthetic MPER peptides partition into membranes and form α-helical structures; peptides corresponding to the gp41 MPER-transmembrane region have been suggested to form trimers in bicelles, but other studies of similar peptides in lipid bilayers or nanodiscs found little evidence of trimers (58–64). These discrepancies may reflect the dynamic nature of the MPER and the dependence of its structure on interactions with both the viral membrane and the rest of the Env ectodomain.

Conserved elements of the HIV-1 gp41 MPER can be targeted by broadly neutralizing antibodies (65). The antibodies generally recognize the MPER in conjunction with the lipid membrane and, in some cases, have been shown to bind a tilted Env and to extract the peptide epitope from the membrane (63, 65–71). Most MPER-directed neutralizing antibodies recognize the CD4-bound Env better than the pretriggered Env, although the 10E8 antibody can bind the functional virion Env spike in the absence of CD4 (72–77). These observations are consistent with the induction of changes in MPER conformation and/or exposure as a result of Env-receptor interactions.

The HIV-1 gp41 MPER has been suggested to play a role in virus entry by promoting the late stages of Env-mediated membrane fusion. Alteration of a conserved tryptophan-rich MPER motif disrupted the ability of Env to expand the initial fusion pore (78, 79). The MPER includes a potential cholesterol-binding motif and some changes in this element diminish Env function (80–84). Finally, functional links between the FPPR and MPER during the formation of the gp41 six-helix bundle have been suggested (85).

Some changes in the gp41 MPER can result in increased HIV-1 sensitivity to neutralization by antibodies directed against gp120 (86–92). The increased neutralization sensitivity resulting from these MPER changes is, in some cases, enhanced by additional gp41 or gp120 changes (86, 90, 91). We hypothesize that the observed increases in neutralization sensitivity result from a loss of MPER-mediated stabilization of the pretriggered (State-1) Env conformation. A corollary of this hypothesis is that the MPER functions to maintain a higher activation barrier between State 1 and downstream conformations of Env. Therefore, Env mutants whose MPER integrity is compromised are more triggerable and predisposed to sampling downstream conformations. A stabilized Env with a larger activation barrier between State 1 and downstream conformations might be expected to resist the disruptive effects of MPER changes. Here, we characterize in detail the effects of MPER changes in the Env of a Tier-2 primary virus, HIV-1_AD8_. We test the above hypothesis by evaluating whether State 1-stabilizing changes in the rest of the Env ectodomain can revert the phenotypes resulting from alteration of the MPER. The phenotypic consequences of cholesterol depletion on HIV-1 infectivity and neutralization sensitivity are compared with those of MPER alteration.

## RESULTS

### Effects of MPER changes on the wild-type HIV-1_AD8_ Env

To evaluate the effects of MPER alterations on the Env of a primary, Tier-2 HIV-1, we individually changed Leu 669, Trp 672, Ile 675, Thr 676 and Leu 679 of the HIV-1_AD8_ Env (Fig. 1A). These MPER residues are more than 98% conserved in HIV-1 strains, with the exception of Thr 676, which naturally tolerates substitutions of serine residues in ∼45% of HIV-1 strains (93). Alteration of the individual MPER residues reduced virus infectivity in all cases except for the T676A change, which supported HIV-1 infection comparably to the wild-type HIV-1_AD8_ Env (Fig. 1B, left panel). To evaluate the sensitivity of the variant Envs to cold, pseudoviruses were incubated on ice for different lengths of time and their infectivity was then measured. Compared to the wild-type HIV-1_AD8_ virus, the mutant viruses exhibited increased sensitivity to cold (Fig. 1B, middle panel). Most of the MPER-modified viruses lost 50% of their infectivity after 4 hours on ice; the T676A virus exhibited a 5-hour half-life on ice, whereas the wild-type HIV-1_AD8_ required 8 hours on ice to lose half of its infectivity. Consistent with its Tier-2 phenotype, the wild-type HIV-1_AD8_ was not neutralized by the poorly neutralizing 19b antibody, which is directed against the gp120 V3 loop (94) (Fig. 1B, right panel). By contrast, the MPER mutant viruses were neutralized by 19b; the T676A virus was less sensitive to 19b neutralization than the other mutant viruses. No additivity or synergy between the MPER changes was observed; double mutants containing L669S and one of the other MPER changes displayed virus sensitivity to cold and 19b neutralization comparable to those of the L669S single mutant (Fig. 1C). Overall, these results support our hypothesis that altered MPER integrity can result in more triggerable Envs that are more prone to making transitions from State 1 to downstream conformations. A more comprehensive and detailed characterization of the effects of MPER changes on the Env ectodomain will be presented below.

**Figure 1.**
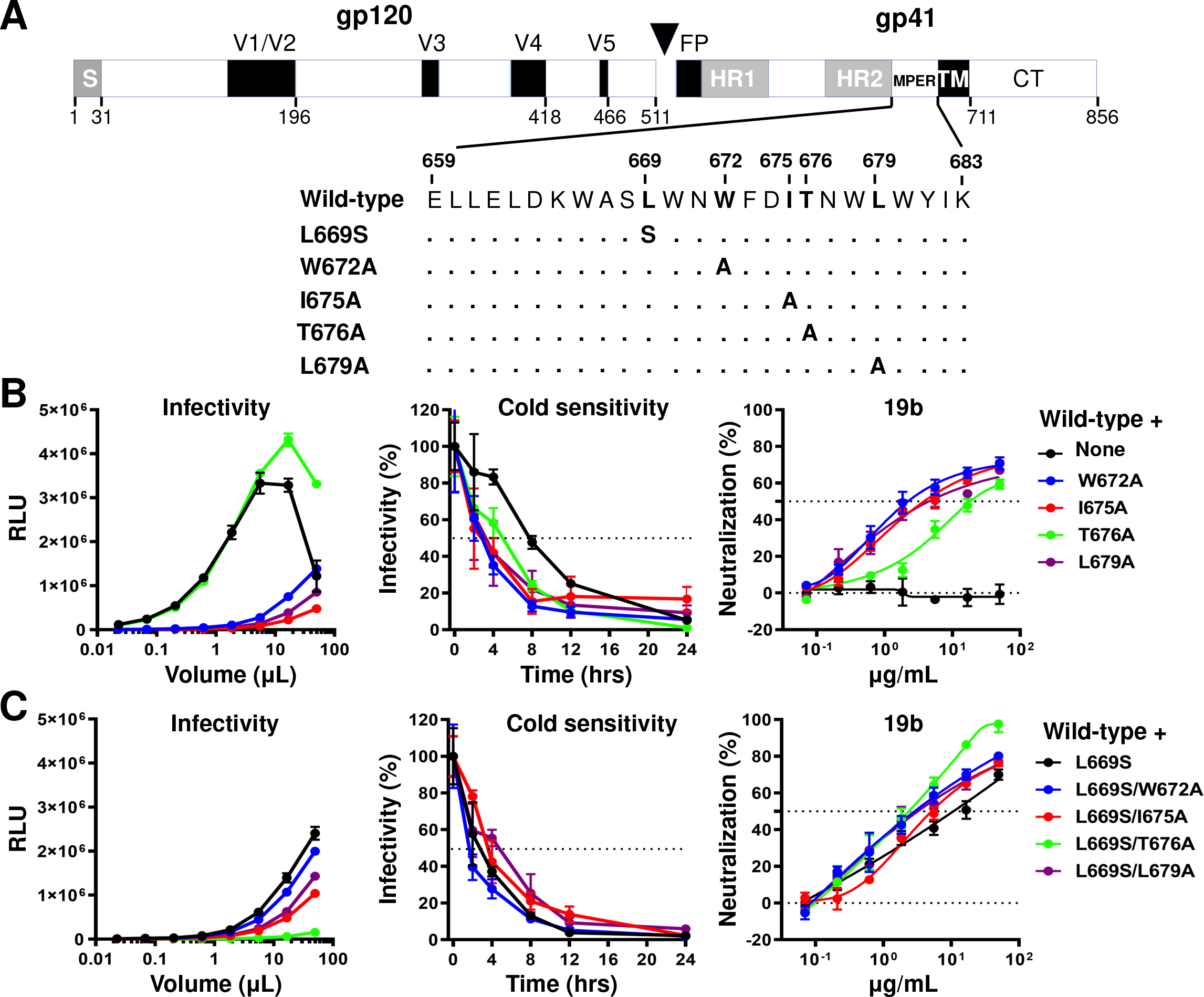
Effects of MPER changes in the wild-type HIV-1_AD8_ Env. (A) A schematic representation of the HIV-1_AD8_ Env glycoprotein is shown, with the gp120-gp41 cleavage site depicted as a black triangle. S, signal peptide; V1-V5, gp120 hypervariable regions; FP, fusion peptide; HR, heptad repeat region; MPER, membrane-proximal external region; TM, transmembrane region; CT, cytoplasmic tail. The MPER amino acid changes studied herein are listed. Standard numbering of HIV-1 Env amino acid residues is used (141). (B, C) HEK293T cells were transfected with pNL4-3.Luc.R-E- and a plasmid expressing the wild-type or MPER-modified HIV-1_AD8_ Env. All Env mutants in C contain the L669S change, either alone or in combination with the other MPER changes. Forty-eight hr later, the cell supernatants were collected, clarified and added to TZM-bl cells in the presence of 20 µg/ml DEAE-Dextran. Forty-eight hr later, the infected cells were lysed, and the luciferase activity (in Relative light units (RLU)) was measured as an indicator of virus infectivity (left panels). In the cold sensitivity assay, viruses were incubated on ice for the indicated times, after which the virus infectivity was measured (middle panels). In the neutralization assay with the 19b poorly neutralizing antibody, viruses were incubated with serial dilutions of 19b for 1 hr at 37°C before TZM-bl cells were added and infectivity was measured. The results for cold sensitivity and 19b neutralization are normalized to those obtained in the absence of virus exposure to cold or 19b antibody. The results shown are representative of those obtained in two independent experiments, expressed as means and standard deviations from triplicate luciferase readings.

### Effects of the MPER changes are minimized in the 2-4 RM6 AE Env

In a previous study, we identified several lysine-rich and well-cleaved HIV-1_AD8_ Env variants with phenotypes consistent with State-1 stabilization (95). Compared to the wild-type HIV-1_AD8_, viruses with these variant Envs were significantly more resistant to cold, soluble CD4, the CD4-mimetic compound BNM-III-170 (96), and to gp120-trimer dissociation in detergent. One such Env variant, 2-4 RM6 AE, was found to tolerate the State 1-destabilizing effects of changes (R542V, I595F and L602H) in or near the gp41 HR1 region (95). Here, we introduced the MPER changes into the 2-4 RM6 AE Env to test whether the phenotypes indicative of State-1 disruption would be less pronounced in the 2-4 RM6 AE Env background than in the context of the wild-type HIV-1_AD8_ Env (Fig. 2A). The MPER mutants exhibited decreases in infectivity relative to the 2-4 RM6 AE parent virus (Fig. 2B, left panel). Strikingly, the MPER-mediated phenotypes associated with State-1 destabilization were minimized or completely reverted in the 2-4 RM6 AE Env background. While the 2-4 RM6 AE viruses with MPER changes were more sensitive to cold relative to the parental 2-4 RM6 AE virus, they were significantly more cold-resistant than wild-type HIV-1_AD8_; the half-lives on ice of the 2-4 RM6 AE MPER mutants were at least 2 days, compared to the 8-hour half-life of wild-type HIV-1_AD8_ (Compare Fig. 2B, middle panel with Fig. 1B, middle panel). Likewise, the MPER modifications in the 2-4 RM6 AE Env did not result in increased sensitivity to neutralization by the 19b anti-V3 antibody (Fig. 2B, right panel). The 2-4 RM6 AE Env with the L669S change tolerated additional MPER changes without effects on cold sensitivity or sensitivity to 19b neutralization (Fig. 2C). These results support the hypothesis that some phenotypic effects of MPER alterations are minimized in the context of an Env that exhibits a more stable pretriggered (State-1) conformation.

**Figure 2.**
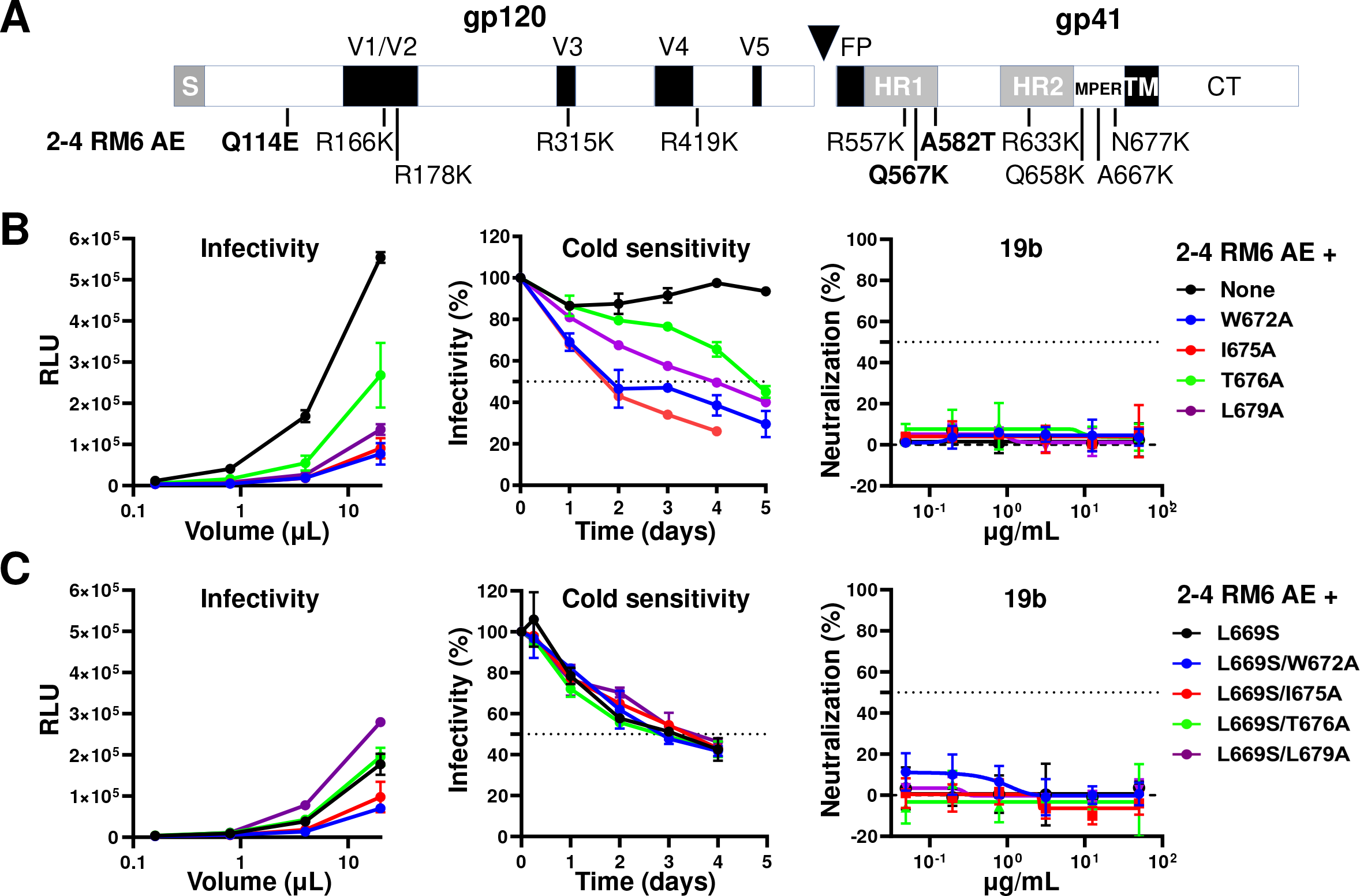
Effects of MPER changes in the State 1-stabilized 2-4 RM6 AE Env. (A) The 2-4 RM6 AE Env is an HIV-1_AD8_ Env variant in which the pretriggered (State-1) conformation is stabilized, compared to the wild-type HIV-1_AD8_ Env (95). Of the 12 changes in the 2-4 RM6 AE Env relative to the wild-type HIV-1_AD8_ Env, the three highlighted changes (Q114E, Q567K and A582T) are the key determinants of the viral phenotypes associated with State-1 stabilization (95). (B, C) The experiments were carried out as described in the legend to Figure 1. Briefly, cell supernatants containing recombinant luciferase-expressing viruses were collected from transiently transfected HEK293T cells, clarified and either used directly for infection (left panels) or incubated with the 19b antibody for 1 hr at 37°C (right panels) before TZM-bl target cells were added. After 48-72 hr, luciferase activity (in RLU) was measured. In the cold sensitivity assay, viruses were incubated on ice for the indicated times before virus infectivity was measured (middle panels). Note the cold resistance of the 2-4 RM6 AE variants, compared with that of wild-type HIV-1_AD8_ in Figure 1B. The results shown are representative of those obtained in two independent experiments, expressed as means and standard deviations from triplicate luciferase readings.

### State 1-stabilizing Env changes lessen the phenotypic effects of MPER alteration

The 2-4 RM6 AE Env differs from the wild-type HIV-1_AD8_ Env by 12 amino acid residues (Fig. 2A). Three substitutions, Q114E, Q567K and A582T, were identified as key determinants of the State 1-stabilizing phenotypes of the 2-4 RM6 AE Env (95). To test our hypothesis that State 1-stabilizing Env changes could revert the State 1-destabilizing effects of MPER changes, we introduced the MPER changes into HIV-1_AD8_ Envs with Q114E/Q567K and Q114E/Q567K/A582T alterations; because the effects of these alterations are additive, the viral phenotypes associated with State-1 stabilization are more robust for the Q114E/Q567K/A582T mutant than for the Q114E/Q567K mutant (95). The MPER changes exerted only modest effects on the infectivity of the Q114E/Q567K and Q114E/Q567K/A582T viruses (Figs. 3A and 3B, left panels). The effects of the MPER changes on virus sensitivity to cold and 19b antibody neutralization were decreased in the Q114E/Q567K background compared with those seen in the context of the wild-type HIV-1_AD8_ (Compare Fig. 3A with Fig. 1B). The half-lives in the cold of the MPER mutants increased from 4-5 hours in the wild-type HIV-1_AD8_ background to around 24 hours (or more for the T676A mutant) in the Q114E/Q567K virus background. Likewise, even at the highest concentrations tested, the 19b antibody only partially inhibited infection of the Q114E/Q567K MPER mutant viruses. The effects of the MPER changes on virus sensitivity to cold and 19b neutralization were further minimized by the addition of the A582T change, a gp41 HR1 substitution that stabilizes the pretriggered (State-1) conformation of Env (97–100) (Fig. 3B). Nearly all of the resistance to the phenotypic effects of MPER changes displayed by the 2-4 RM6 AE Env could be accounted for by the Q114E/Q567K/A582T changes (Compare Fig. 3B with Fig. 2B). These results support the hypothesis that phenotypic effects of MPER alterations can be reverted by State 1-stabilizing changes in the Env ectodomain outside of the MPER.

**Figure 3.**
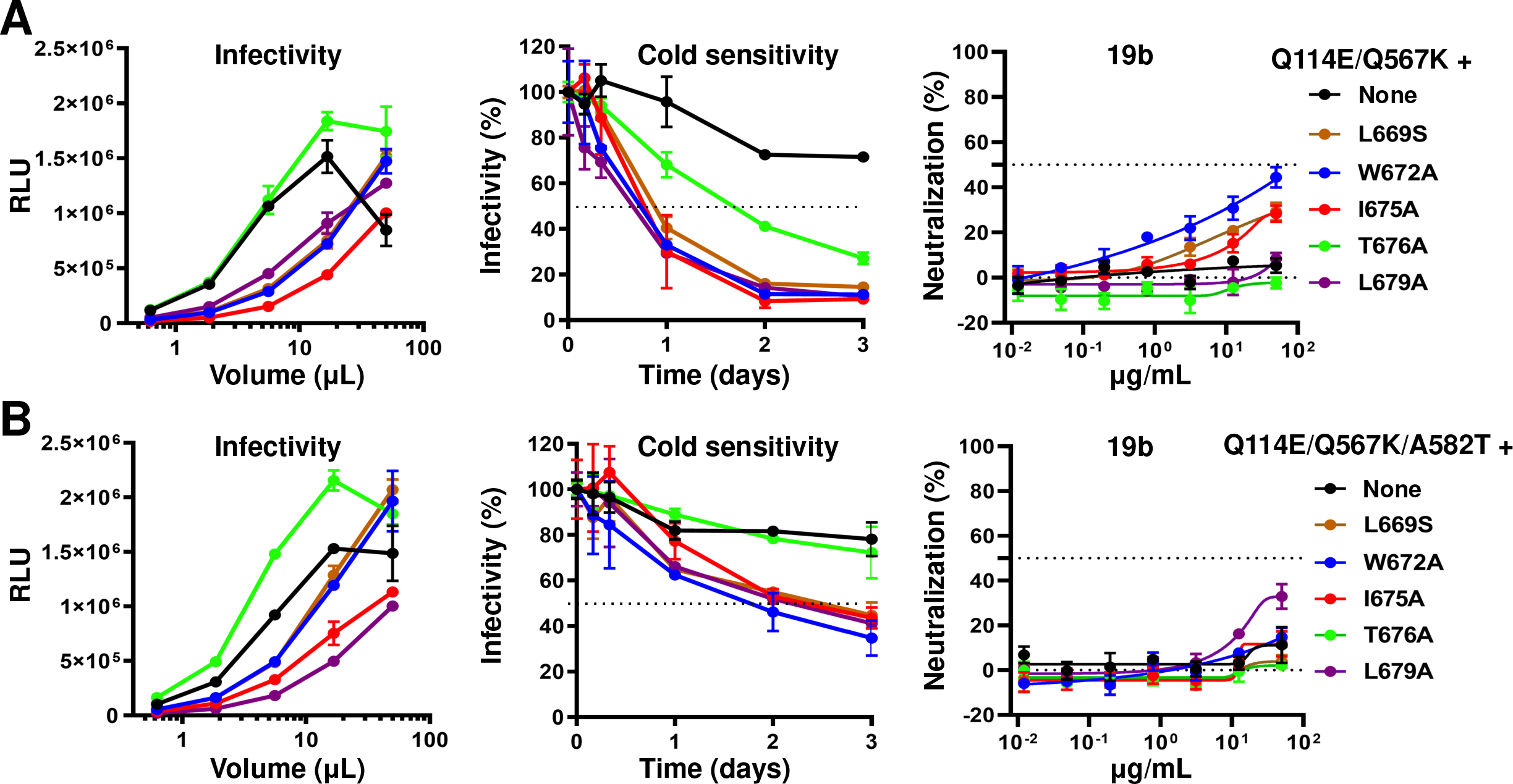
Effects of MPER changes in HIV-1_AD8_ Envs with State 1-stabilizing changes. The indicated MPER changes were introduced into Envs containing Q114E/Q567K (A) or Q114E/Q567K/A582T substitutions (B) that have previously been shown to stabilize a pretriggered (State-1) conformation (95). The experiments were carried out as described in the legend to Figure 1. Briefly, recombinant viruses produced from HEK293T cells transfected with pNL4-3.Luc.R-E- and an Env-expressing plasmid were incubated with TZM-bl target cells for 48 hr, after which luciferase activity was measured (left panels). In the cold sensitivity assay, virus was incubated on ice before addition to target cells (middle panels). In the neutralization assay, virus was preincubated with the 19b antibody for 1 hr at 37°C before TZM-bl cells were added to the mixture (right panels). The results shown are representative of those obtained in two independent experiments, expressed as means and standard deviations from triplicate luciferase readings.

### Overall effects of the MPER changes in different Env backgrounds

The above findings suggest that MPER changes can promote Env transitions from its pretriggered conformation to downstream states. Furthermore, Envs exhibiting phenotypes indicative of increased State-1 stability are less susceptible to some of the consequences of MPER alteration. To obtain a more complete picture, we evaluated the range of Env phenotypes that potentially might be affected by MPER modification. We compared the MPER mutant phenotypes in the backgrounds of the wild-type HIV-1_AD8_ Env and the State 1-stabilized Envs.

The non-covalent association of gp120 with detergent-solubilized Env trimers is prone to disruption, providing information about the stability of the Env complex. The gp120-trimer association was evaluated by a Ni-NTA precipitation assay (101). For this assay, HOS cells transiently expressing Envs with C-terminal His_6_ tags were lysed in buffers containing the Cymal-5 detergent. The clarified lysates were incubated with Ni-NTA beads, and the precipitated Envs (as well as an “Input” sample from the cell lysates obtained prior to incubation with the Ni-NTA beads) were Western blotted. The results of the assays conducted on the Env variants at 25°C and 4°C are shown in Figure 4. Comparison of the gp120 and gp160 bands in the “Input” samples indicates that the MPER changes did not detectably affect either the proteolytic processing or the subunit association of the Env variants. As previously seen (95), the processing of the 2-4 RM6 AE, Q114E/Q567K and Q114E/Q567K/A582T Envs was at least as efficient as that of the wild-type HIV-1_AD8_ Env. The gp120-trimer association in detergent lysates is reflected in the gp120:gp160 ratio in the Ni-NTA precipitates compared with the gp120:gp160 ratio in the Input samples. As previously seen (101), the gp120-trimer association of the wild-type HIV-1_AD8_ Env in detergent lysates was lower at 25°C than at 4°C. Also consistent with a previous study (95), the Env variants with the State 1-stabilizing modifications (2-4 RM6 AE, Q114E/Q567K and Q114E/Q567K/A582T) displayed stronger gp120-trimer association in detergent lysates than the wild-type HIV-1_AD8_ Env. The MPER changes did not significantly affect gp120-trimer association in any of the Env backgrounds. Neither did the MPER changes in the wild-type HIV-1_AD8_ background affect spontaneous gp120 shedding from cell-surface Envs or Env incorporation into virus particles (data not shown). These results indicate that the general structural integrity of the Env trimer is maintained in the MPER mutants.

**Figure 4.**
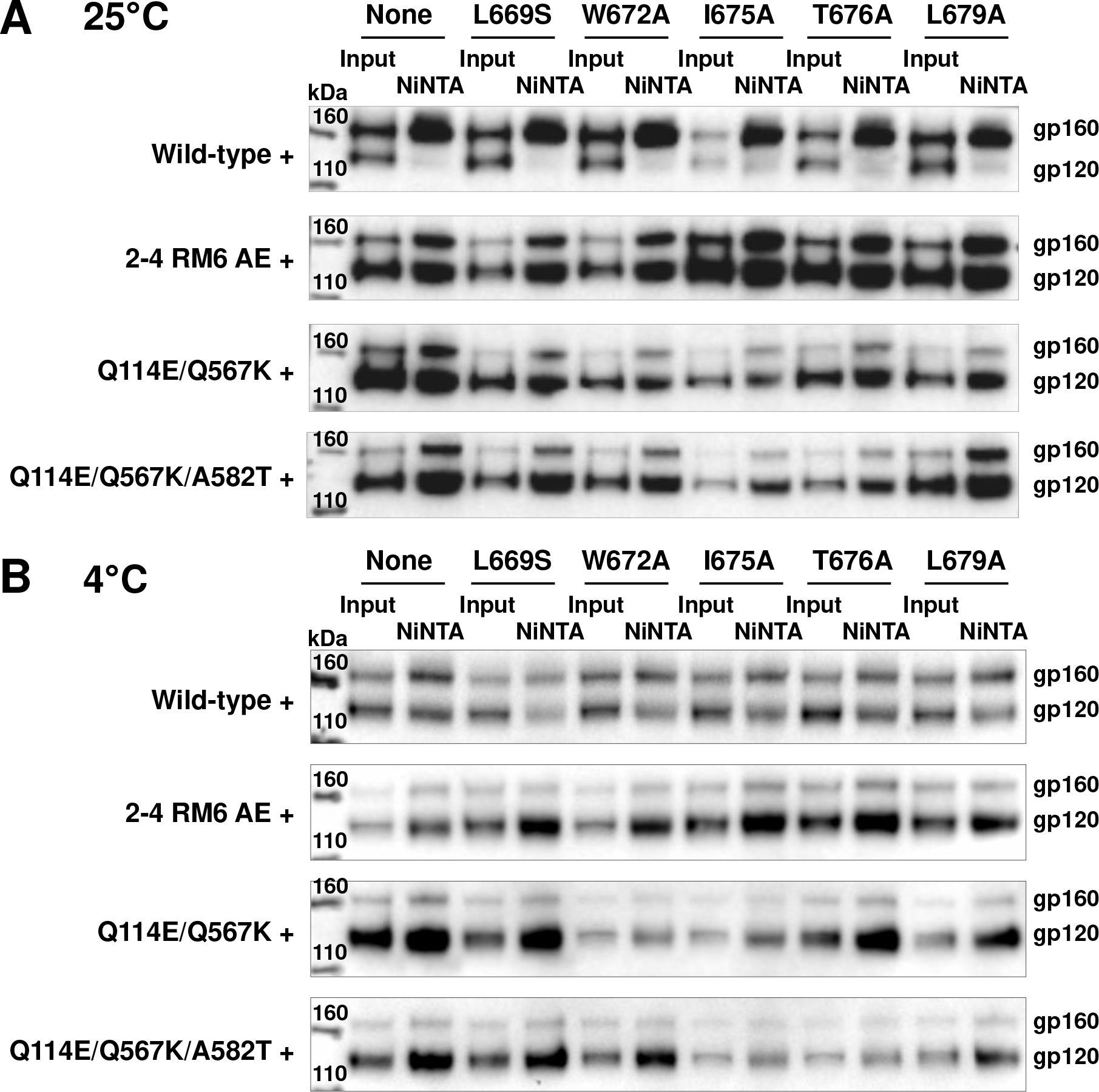
Effects of the MPER changes on gp120-trimer association in detergent. HOS cells transiently expressing Env glycoproteins tagged at the C-terminus with His_6_ were lysed and incubated with Ni-NTA beads for 1 hr at 25°C (A) or at 4°C (B). Total cell lysates (Input) and Ni-NTA precipitates were Western blotted with a goat anti-gp120 antibody. The results shown are representative of those obtained in two independent experiments.

To examine more thoroughly the effects of the MPER alterations on the conformation of the functional Envs, we carried out virus neutralization assays using monoclonal antibodies and inhibitors targeting different gp120 and gp41 regions (Fig. 5). Poorly neutralizing antibodies directed against gp120 included F105 against the CD4 binding site (CD4bs) (102), 17b against a CD4-induced (CD4i) epitope (103) and 19b against the V3 region (94). The gp120-directed broadly neutralizing antibodies included VRC03 against the CD4bs (104), PG9 and PGT145 against quaternary V2 epitopes (105, 106) and PGT151 against the gp120-gp41 interface (107). Virus entry inhibitors directed against gp120 included soluble CD4-Ig (sCD4-Ig), the small-molecule CD4-mimetic compound (CD4mc) BNM-III-170 (96) and the State 1-preferring small molecule, BMS-806 (108). The gp41-directed ligands included the T20 peptide, which mimics the HR2 region and targets the HR1 coiled coil (109), and the broadly neutralizing 10E8 antibody against the MPER (60, 68, 77).

**Figure 5.**
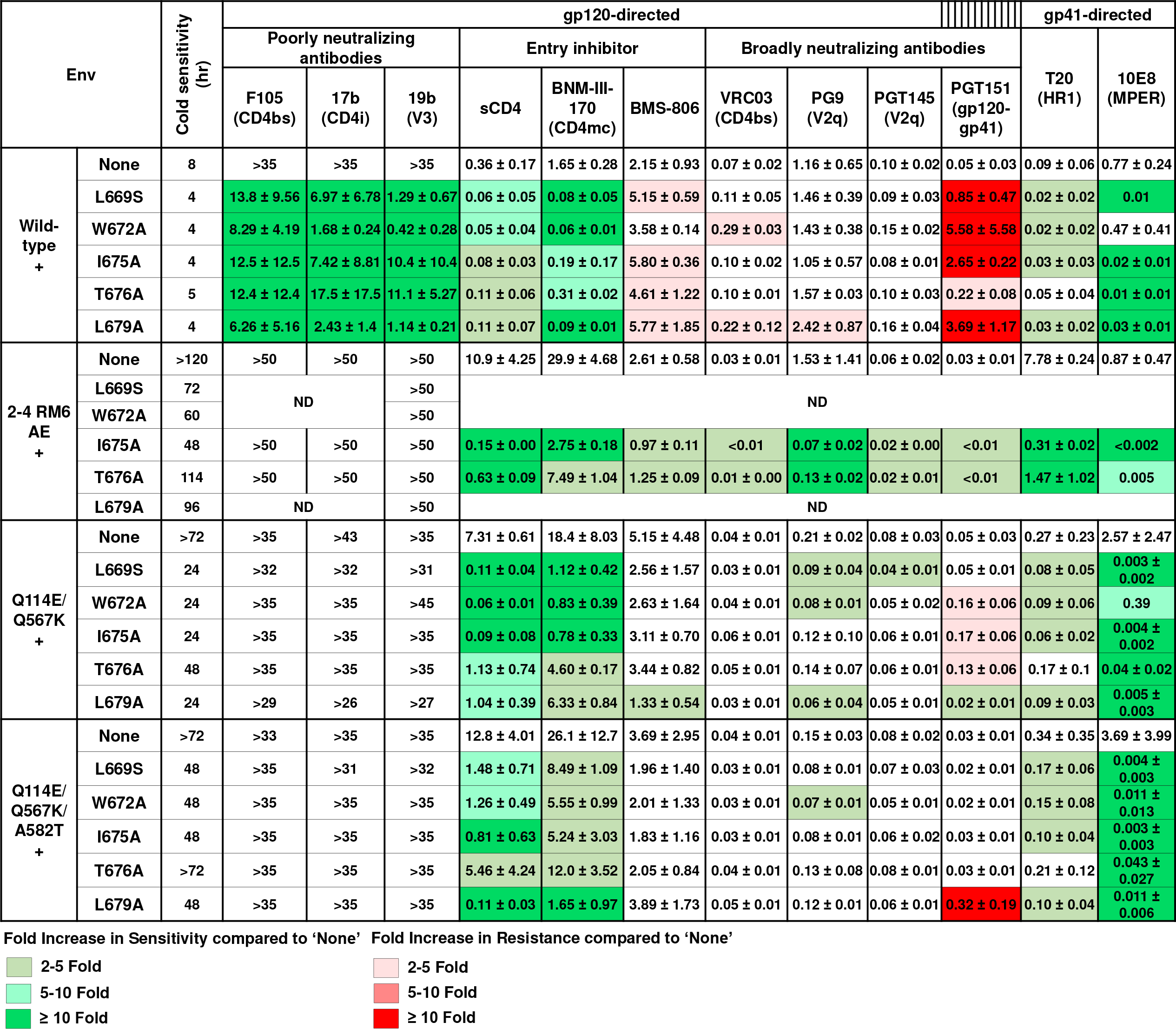
Phenotypes of viruses with Envs containing MPER changes. Pseudoviruses containing the indicated Env variants were tested for sensitivity to cold and to antibodies/compounds directed against gp120 or gp41. Poorly neutralizing antibodies and broadly neutralizing antibodies are indicated. The 50% inhibitory concentrations (IC_50_ values) of the Env ligands are reported in µg/ml except for BNM-III-170 (in µM) and BMS-806 (in nM). Some of the phenotypes were not determined (ND) due to the low infectivity of the viruses. Values are colored based on their fold increase in sensitivity (green) or resistance (red) compared to the relevant parental virus (labeled “None”). The parental and associated MPER mutant viruses were compared in parallel experiments. The results shown are means and standard deviations from two independent experiments. Additional experiments validated the relative IC_50_ values of the parental viruses for sCD4-Ig, BNM-III-170, BMS-806 and cold. The mean IC_50_ values of T20 and 10E8.v4 were obtained in a side-by-side comparison of the four parental viruses (wild-type HIV-1_AD8_, 2-4 RM6 AE, Q114E/Q567K and Q114E/Q567K/A582T): 0.56, 0.64, 2.5 and 0.89 µg/ml of T20, respectively; 1.7, 0.11, 2.4 and 2.0 µg/ml of 10E8.v4, respectively.

The phenotypes resulting from the MPER changes depended upon the Env background, as shown in Fig. 5 and summarized schematically in Fig. 6. In the background of the wild-type HIV-1_AD8_ Env, the MPER mutants were more sensitive to neutralization by the poorly neutralizing F105, 17b and 19b antibodies, and by sCD4 and the CD4mc BNM-III-170. These MPER mutants also exhibited modest increases in sensitivity to T20 compared with the wild-type HIV-1_AD8_ parent. Along with increased cold sensitivity, these MPER mutant phenotypes are consistent with destabilization of the pretriggered (State-1) conformation. Relative to wild type HIV-1_AD8_, the MPER mutants were significantly more resistant to the broadly neutralizing PGT151 antibody; some MPER mutants also exhibited slight resistance to VRC03 and BMS-806, two State-1 preferring Env ligands (13, 110–112). No significant changes were detected in the sensitivity of the MPER mutants to neutralization by the PG9 and PGT145 antibodies against V2 quaternary epitopes. The MPER mutants were more sensitive to neutralization by the 10E8 antibody, except for W672A; the W672A change affects the MPER epitope for this antibody (77). In summary, multiple single-residue changes in the MPER of the wild-type HIV-1_AD8_ Env led to phenotypes indicative of global disruption of the pretriggered (State-1) conformation (Compare panels 1 and 1b in Fig. 6).

**Figure 6.**
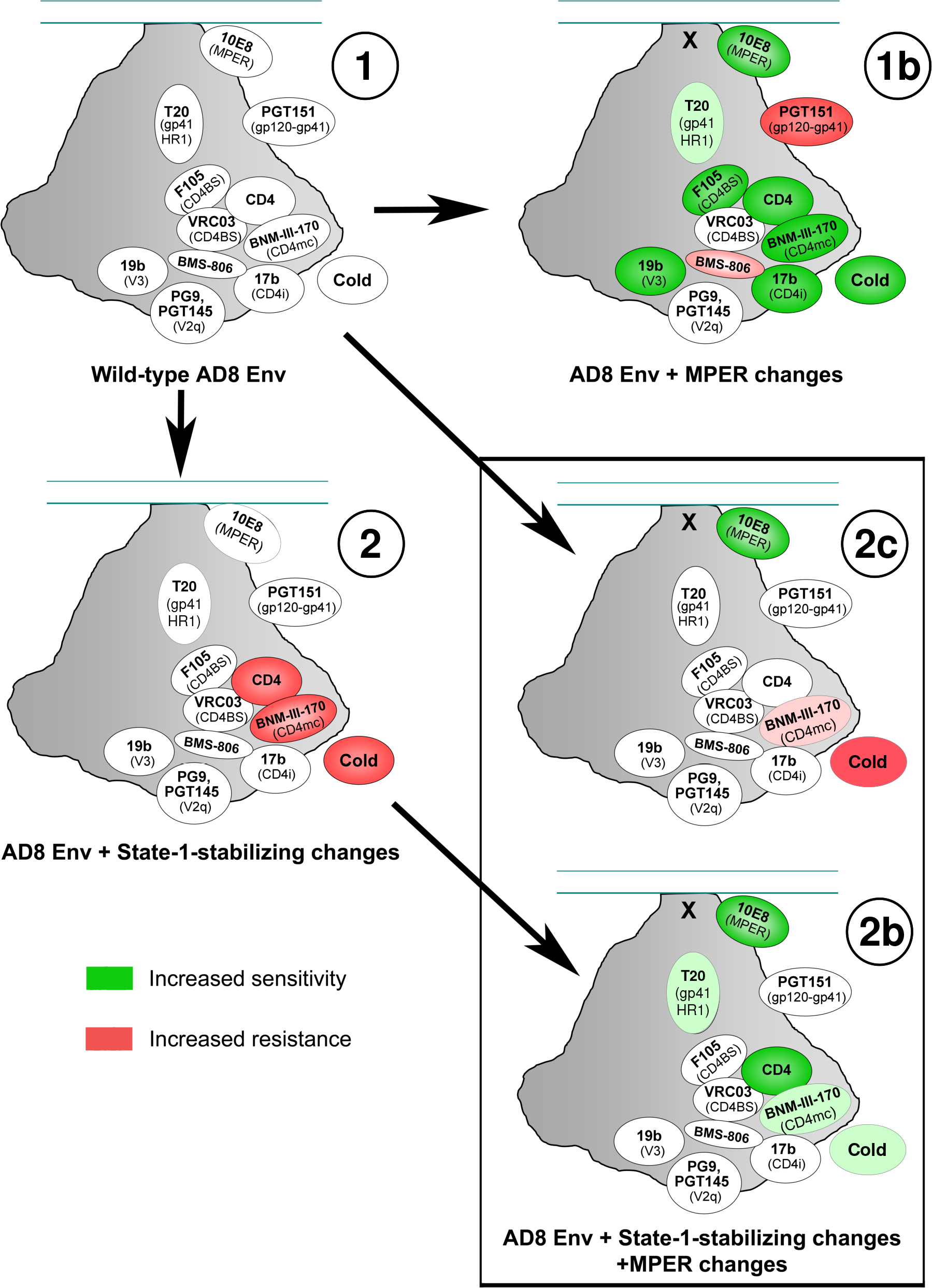
Summary of the phenotypic effects of the MPER and State 1-stabilizing changes on virus sensitivity to Env ligands or cold exposure. The overall phenotypic effects of MPER changes and State 1-stabilizing changes in Env on virus sensitivity to Env ligands and cold exposure (from Figure 5 and additional side-by-side experiments comparing the four parental Envs) are summarized here. On a schematic view of the HIV-1 Env trimer, the approximately positioned ligands are colored according to increased sensitivity (green) or increased resistance (red); in each case, the phenotypes of the Env variant following the arrow are compared with those of the Env variant preceding the arrow. For example, the phenotypes of wild-type HIV-1_AD8_ Envs without and with MPER changes can be compared in panels 1 and 1b, respectively. For the depictions of the State 1-stabilized Envs, the data from the Q114E/Q567K and Q114E/Q567K/A582T parental Envs were used, to avoid potential effects of the other changes in the 2-4 RM6 AE Env unrelated to State-1 stabilization. Examination of panels 1b and 2b allows a comparison of the effects of the MPER changes in the context of the wild-type HIV-1_AD8_ Env and the State 1-stabilized Envs, respectively. The phenotypes of the wild-type HIV-1_AD8_ Env, the HIV-1_AD8_ Env with MPER changes, and the State 1-stabilized Envs with MPER changes can be compared in panels 1, 1b and 2c, respectively.

As expected (95), compared to the wild-type HIV-1_AD8_, viruses with the 2-4 RM6 AE, Q114E/Q567K and Q114E/Q567K/A582T Envs were significantly resistant to cold, sCD4-Ig and BNM-III-170 (Fig. 5), phenotypes consistent with State-1 stabilization. The IC_50_ values of the other Env ligands tested are not directly comparable for these four parental Envs (rows labeled “None” in Fig. 5), as they were not tested in parallel. However, the relative sensitivity of viruses pseudotyped with the wild-type HIV-1_AD8_, Q114E/Q567K and Q114E/Q567K/A582T Envs to these other ligands was evaluated in separate assays and found not to differ significantly, as summarized in panels 1 and 2 of Fig. 6. The relative resistance of the Q114E/Q567K and Q114E/Q567K/A582T pseudoviruses specifically to sCD4-Ig, CD4mc and cold is indicative of the higher activation barrier of these Envs to CD4-induced transitions from State 1 to downstream conformations. When the MPER changes were introduced into these Env backgrounds, two major differences in the resulting viral phenotypes were observed compared to those seen in the wild-type HIV-1_AD8_ background. First, the MPER mutants remained resistant to poorly neutralizing antibodies. Second, relative to the parent viruses, the MPER mutants generally did not display resistance to PGT151, VRC03 or BMS-806 (Compare panels 2 and 2b in Fig. 6). In all Env backgrounds, relative to the parent virus, alteration of the MPER led to enhanced viral sensitivity to cold, sCD4-Ig, BNM-III-170, T20 and 10E8 (Compare panels 1 and 1b and panels 2 and 2b in Fig. 6). In contrast to the globally “opened” Env resulting from the introduction of MPER changes into the wild-type HIV-1_AD8_ background (panel 1b in Fig. 6), viruses with both MPER and State 1-stabilizing Env changes differ minimally from wild-type HIV-1_AD8_ (Compare panels 1 and 2c in Fig. 6). The viruses with combined MPER and State 1-stabilizing Env changes displayed an increased sensitivity to 10E8 neutralization, possibly reflecting local effects on the MPER, but exhibited none of the phenotypes associated with open, State 1-destabilized Envs.

The above findings suggest that Envs with MPER alterations are more prone to make transitions from the pretriggered (State-1) conformation to more open downstream conformations. We evaluated whether infection by wild-type and MPER mutant viruses was activated with different levels of efficiency by sCD4-Ig and the CD4mc, BNM-III-170. In this assay, viruses were spinoculated on CD4-negative, CCR5-expressing Cf2Th-CCR5 cells, after which sCD4-Ig or BNM-III-170 was added. Forty-eight hours later, virus infection was measured. In the absence of sCD4-Ig or BNM-III-170, a very low background was observed in this assay. Wild-type HIV-1_AD8_ infection of the Cf2Th-CCR5 cells was significantly enhanced by sCD4-Ig and BNM-III-170 (Fig. 7A). Infection by the T676A mutant virus was activated at lower doses of sCD4-Ig and BNM-III-170 than wild-type HIV-1_AD8_ infection. Infection by the other MPER mutant viruses was not detectably enhanced by sCD4-Ig or BNM-III-170, perhaps due to their intrinsically low infectivity. Infection by viruses with the 2-4 RM6 AE Env was activated only at the highest BNM-III-170 concentration tested, consistent with the lower expected triggerability of this State 1-stabilized variant (95). Infection by viruses with MPER-altered 2-4 RM6 AE Envs was not detectably activated by either sCD4-Ig or BNM-III-170. These observations are consistent with the T676A change in the MPER increasing the responsiveness of the HIV-1_AD8_ Env to CD4 and a CD4mc.

**Figure 7.**
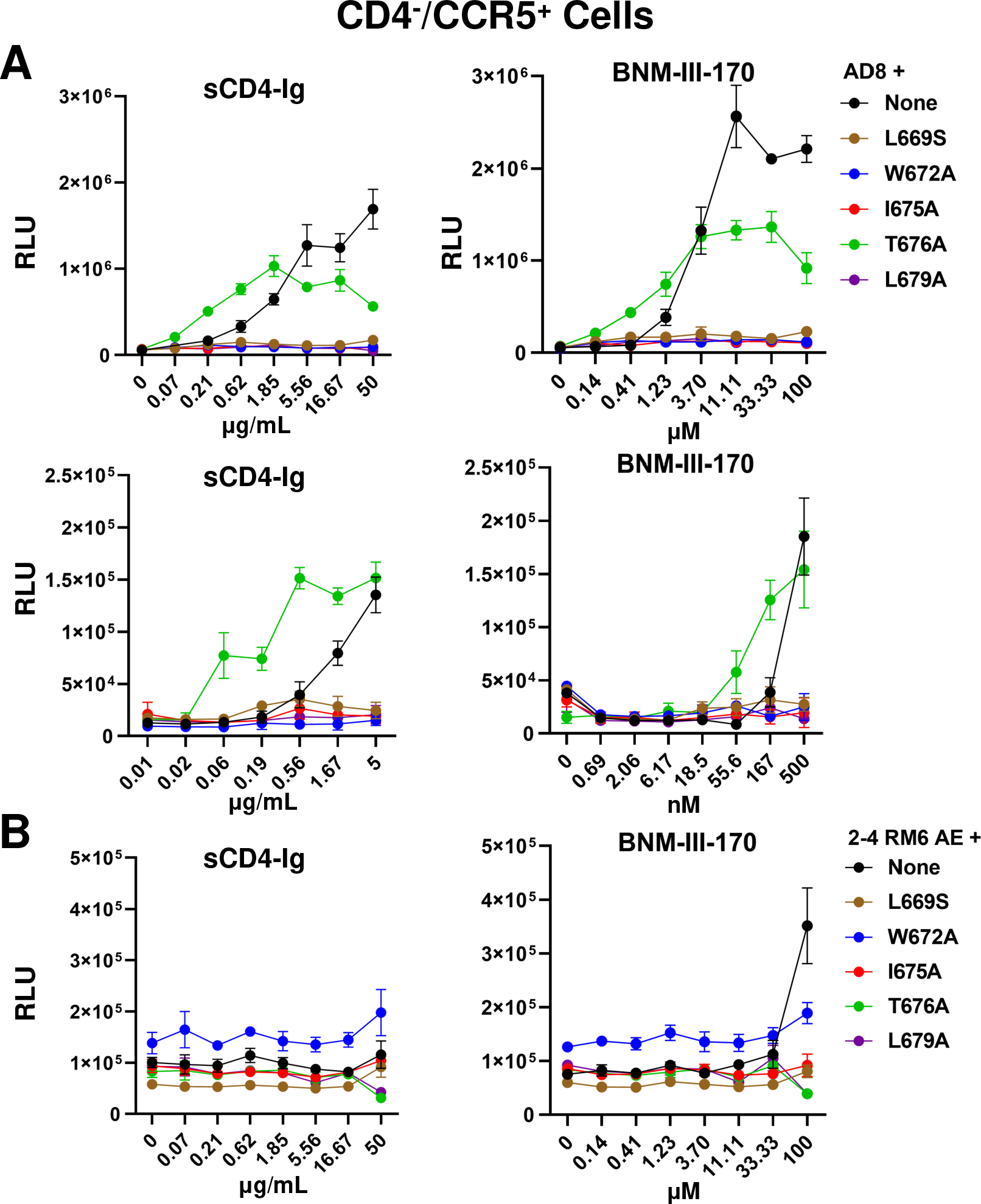
CD4- and CD4mc-mediated activation of infection of CD4-negative cells by HIV-1 Env variants. Wild-type HIV-1_AD8_ (A) and 2-4 RM6 AE viruses (B) containing the indicated MPER changes were spinoculated on CD4-negative, CCR5-expressing Cf2Th-CCR5 cells, after which serial dilutions of sCD4-Ig or BNM-III-170, a CD4-mimetic compound, were added. Note that a lower range of sCD4-Ig and BNM-III-170 was used in the lower panels of A. Forty-eight hr later, the cells were lysed and luciferase activity was measured. The results shown are representative of those obtained in two independent experiments, expressed as means and standard deviations from triplicate luciferase readings.

### Effect of membrane cholesterol on HIV-1 Env phenotypes

The MPER contains a conserved cholesterol recognition motif spanning residues 679-683 (80–83). Changes in this motif can result in viruses with impaired replicative ability and increased sensitivity to cold, poorly neutralizing antibodies and CD4mc (84, 87, 88, 92). Depletion of cholesterol from the virus particles of certain HIV-1 strains produces similar effects (92, 113–118). We asked whether changes that stabilize the State-1 conformation would alter virus susceptibility to decreases in membrane cholesterol. Methyl-β-cyclodextrin (MBCD) was used to deplete cholesterol from the viral membrane (119). The wild-type HIV-1_AD8_, Q114E/Q567K, Q114E/Q567K/A582T and 2-4 RM6 AE viruses were all inhibited comparably by MBCD (data not shown). These results indicate that stabilization of the pretriggered (State-1) Env conformation does not bypass or diminish the requirement of HIV-1 for cholesterol in the membrane.

Next, we asked whether cholesterol depletion would result in State 1-destabilized phenotypes similar to those resulting from MPER changes. The viruses were incubated with MBCD at a 50% inhibitory concentration for one hour and then tested for sensitivity to neutralization by antibodies and BNM-III-170. As seen in Fig. 8, no difference was observed in the sensitivity of wild-type HIV-1_AD8_, Q114E/Q567K, Q114E/Q567K/A582T and 2-4 RM6 AE viruses to poorly neutralizing antibodies (19b, 17b, F105), the broadly neutralizing antibody PGT151, or the CD4mc BNM-III-170. However, the MBCD-treated viruses were slightly more sensitive than the untreated control viruses to neutralization by 10E8.v4, an optimized version of the 10E8 antibody against the MPER (120). This observation verifies that viral components near or within the membrane were affected by MBCD treatment. Some local effects of cholesterol removal from the viral membrane may resemble those resulting from MPER changes, which also result in increased sensitivity to 10E8 antibody neutralization (See Figs. 5 and 6 above). However, the more global effects of MPER changes on HIV-1 Env conformation are distinct from the effects of cholesterol removal from the viral membrane.

**Figure 8.**
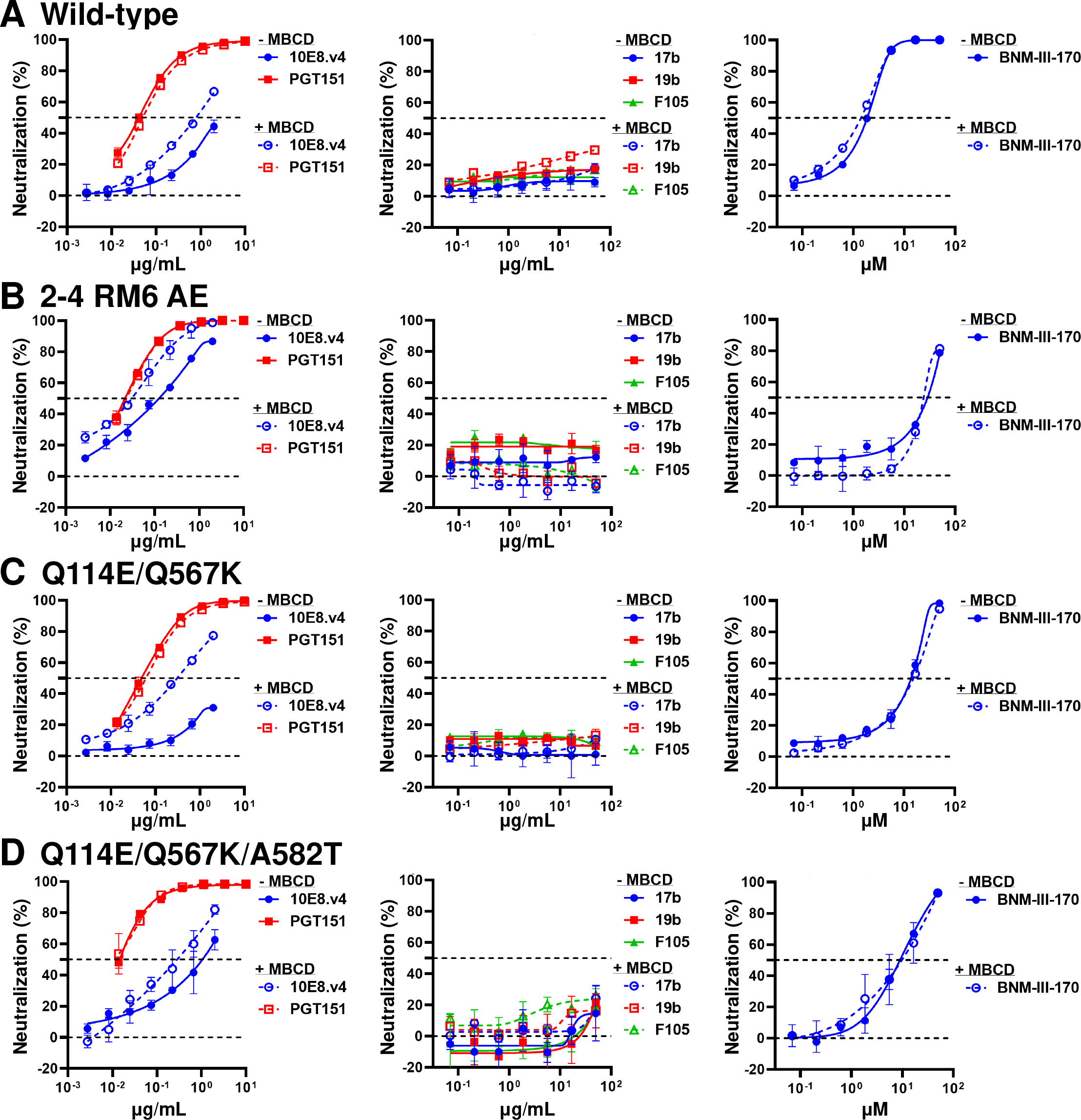
Effects of cholesterol depletion on virus sensitivity to neutralization. The wild-type HIV-1_AD8_ (A), 2-4 RM6 AE (B), Q114E/Q567K (C) and Q114E/Q567K/A582T (D) viruses were incubated with methyl-β-cyclodextrin (MBCD) at the pre-determined IC_50_ concentration for each virus for 1 hr at 37°C. Then the indicated antibodies or BNM-III-170 were added and the mixture was incubated for another hour. The mixture was added to TZM-bl target cells and 48 hr later, luciferase activity was measured. The results shown are representative of those obtained in two independent experiments, expressed as means and standard deviations from triplicate luciferase readings.

## DISCUSSION

Changes introduced into several highly conserved amino acid residues in the HIV-1 gp41 MPER resulted in remarkably similar viral phenotypes. This observation suggests that each of these changes exerted a significant impact on MPER integrity. Only the T676A mutant, affecting a residue that naturally tolerates Thr/Ser polymorphism (93), exhibited phenotypes intermediate between those of the parental viruses and the other mutants. Despite efficient processing and subunit association, the MPER mutants exhibited modest decreases in infectivity, in keeping with the role proposed for this gp41 region in membrane fusion and virus entry (78–85). Our results highlight another key role played by the gp41 MPER, namely, the regulation of the conformation of the HIV-1 Env ectodomain. MPER alteration led to both local and distant conformational effects. In gp41, MPER changes resulted in enhanced sensitivity to the 10E8 antibody against the MPER and to the T20 peptide against HR1 (Fig. 6, panels 1 and 1b), as has been previously seen (121). The increased sensitivity of some MPER mutants to other MPER-directed antibodies, 2F5 and 4E10, has been suggested to result from enhanced/prolonged exposure of the MPER on virus particles (87-89, 122, 123). At a distance, in gp120, the MPER changes resulted in increased sensitivity to sCD4-Ig, a CD4-mimetic compound, BNM-III-170, and several poorly neutralizing antibodies. This pattern of enhanced virus sensitivity to ligands that prefer State-2/3 Env conformations, along with increased virus inactivation in the cold, is strongly associated with destabilization of the pretriggered (State-1) Env conformation (86-92, 95, 110, 124-139). Infection of CD4-negative, CCR5-expressing cells by the T676A mutant, which retains the highest level of infectivity among the MPER mutants, was also more easily activated by sCD4-Ig and BNM-III-170 than the wild-type HIV-1_AD8_. Together, these findings support the hypothesis that the MPER changes destabilize State 1, predisposing Env to assume downstream conformations.

To test this hypothesis, we examined whether the viral phenotypes associated with the MPER changes would be mitigated in the context of State 1-stabilized Envs. These previously reported viruses with State 1-stabilized Envs retained the resistance to the poorly neutralizing antibodies seen for the Tier-2 parental virus, HIV-1_AD8_, but were relatively resistant to CD4-Ig, the CD4mc BNM-III-170, and exposure to cold (Fig. 6, panels 1 and 2) (95). Remarkably, in the context of State 1-stabilized Envs, the multiple phenotypic effects of MPER changes on wild-type HIV-1_AD8_ were nullified, except for an increased sensitivity to the 10E8 antibody against the MPER (Compare panels 1b and 2c in Fig. 6). Apparently, the State 1-destabilizing effects of the MPER changes can be balanced by the State 1-stabilizing changes present in the 2-4 RM6 AE, Q114E/Q567K and Q114E/Q567K/A582T Envs. This mechanism may underlie the HIV-1 strain dependence of the phenotypes observed for some MPER changes (89, 92), predicting lower penetrance (less apparent phenotypes) in the context of Envs with lower triggerability (more stabilized State-1 conformations). For example, introduction of the L669S change into the Tier-2 HIV-1_AD8_ Env in our study generated a virus that exhibited increased sensitivity to multiple poorly neutralizing antibodies (Fig. 6, panels 1 and 1b). By contrast, the same L669S change in a Tier-3 HIV-1_253-11_ Env resulted in more than 10-fold increases in sensitivity to the MPER antibodies 2F5, 4E10 and 10E8, but no differences in sensitivity to the F105, 17b and PGT151 antibodies (89). In the latter instance, the phenotypic effects of the L669S MPER change may have been offset by the low triggerability and strong State-1 stability of the Tier-3 HIV-1 Env.

The mechanism by which the MPER contributes to the stability of the State-1 Env conformation is unknown, but potentially involves direct interactions between the MPER and the viral membrane and/or other gp41 elements, as well as indirect effects on gp120. Based on studies of MPER peptides, the MPER may be partly submerged in the lipid environment of the viral membrane (58–64). Such membrane interactions could stabilize the position and orientation of the Env spike, influencing trimer symmetry and packing of the Env protomers. Changes in MPER conformation may be allosterically coupled to entry-related conformational changes in the rest of the Env ectodomain. Indeed, destabilization of the pretriggered (State-1) Env conformation by CD4 binding or alteration of “restraining residues” in gp120 can lead to increased MPER exposure and enhanced HIV-1 sensitivity to neutralization by MPER-directed antibodies (72-77, 110, 134). Such allosteric coupling between gp120 and gp41 conformations has been suggested by smFRET studies (111), and helps explain the observed synergy between gp120 and gp41 changes in allowing HIV-1 to become CD4-independent (86).

The effects of cholesterol removal from the viral membrane mediated by MBCD differed from the effects of MPER changes. As expected (113–118), MBCD treatment resulted in a decrease in HIV-1 infectivity; however, the apparent requirement of HIV-1 infectivity for membrane cholesterol was not bypassed or diminished by the State 1-stabilizing Env modifications. At MBCD concentrations that reduced HIV-1 infectivity by 50%, global effects on virus sensitivity to neutralizing antibodies were not detected. However, increased sensitivity to 10E8 antibody neutralization was observed, suggesting that cholesterol removal from the viral membrane can increase the efficacy of an anti-MPER antibody. It has been proposed that the 10E8 antibody extracts its peptide epitope from the viral membrane, a process that may be facilitated in a membrane lacking cholesterol (60, 62, 63, 68, 71). In summary, although local effects of MBCD on the MPER were observed, the global effects of MPER changes on HIV-1 Env conformation apparently are distinct from the effects of cholesterol removal from the viral membranes.

The sensitivity of the State-1 HIV-1 Env conformation to MPER disruption has implications for structural and vaccine studies. In most preparations of soluble or detergent-solubilized HIV-1 Env trimers, the MPER has been deleted or is disordered (24, 25, 47–53, 107). Single-molecule FRET studies indicate that the State-1 conformation is not well represented in these Env preparations (111, 140). Thus, detailed information about the State-1 Env conformation, the major target for broadly neutralizing antibodies (13-15, 18, 111, 112), is currently unavailable. Remedying this gap in knowledge may require approaches to maintain MPER integrity in Env preparations or, alternatively, minimizing the impact of MPER disruption on Env conformation. The State 1-stabilized Env designs described here offset many of the effects of MPER changes on functional Env conformation. These Env variants could assist efforts to characterize the elusive State-1 conformation and to maintain this native conformation in vaccine immunogens.

## MATERIALS AND METHODS

### Cell lines

HEK293T, TZM-bl and HOS (ATCC) were cultured in Dulbecco modified Eagle medium (DMEM) supplemented with 10% fetal bovine serum (FBS) and 100 µg/ml penicillin-streptomycin (Life Technologies). CD4 negative, CCR5-expressing Cf2Th-CCR5 cells were cultured in the same medium supplemented with 400 µg/ml G418. Expi293 cells (Invitrogen) were maintained in suspension culture in Expi293TM Expression Medium, supplemented with 50 units/ml penicillin and 50 µg/ml streptomycin.

### Antibodies and sCD4-Ig

Poorly neutralizing antibodies (F105, 17b and 19b) and broadly neutralizing antibodies (VRC03, PG9, PGT145, PGT151, 10E8 and 10E8.v4) against the HIV-1 Env were used in this study. Antibodies were produced by co-transfecting paired heavy- and light-chain antibody-expressing plasmids at a ratio of 1:1 using FectoPRO DNA Transfection Reagent (VWR) into Expi293 cells. Antibodies were purified from the cell medium 5 days after transfection by affinity chromatography using Protein A columns (Thermo Fisher Scientific). Soluble CD4-Ig (sCD4-Ig) encoding the first two domains of human CD4 fused to an antibody Fc was expressed and purified as described above for the antibodies.

### Plasmids

The wild-type HIV-1_AD8_, 2-4 RM6 AE, Q114E/Q567K and Q114E/Q567K/A582T Envs were cloned into the pSVIIIenv expression plasmid, using the Kpn I and BamHI sites, as described previously (95). The signal peptide/N-terminus (residues 1-33) and the cytoplasmic tail C-terminus (residues 751-856) of these Envs are derived from the HIV-1_HXBc2_ Env (86). All the Envs contain a His_6_ tag (GGHHHHHH or GGGHHHHHH) at the carboxyl terminus. The MPER changes (L669S, W672A, I675A, T676A and L679A) were introduced using the QuikChange Lightning Site-Directed Mutagenesis Kit (Agilent Technologies). All the mutations were confirmed by DNA sequencing. Antibody-expressing plasmids were obtained from Dennis Burton (Scripps), Peter Kwong and John Mascola (NIH Vaccine Research Center), Barton Haynes (Duke University), Hermann Katinger (Polymun), James Robinson (Tulane University), Marshall Posner (Mount Sinai Medical Center), David Ho (Columbia University Vagelos College of Physicians and Surgeons) and the NIH HIV Reagent Program.

### Virus infectivity and cold sensitivity

Pseudoviruses bearing the wild-type or variant HIV-1 Envs were produced by co-transfecting the Env-expressing pSVIIIenv plasmid and the luciferase-encoding pNL4-3.Luc.R-E-vector (NIH HIV Reagent Program) at a 1:1 ratio into HEK293T cells using polyethylenimine (Polysciences). The cell supernatants containing pseudoviruses were harvested 48 hr later, clarified by low-speed centrifugation (2000 rpm for 10 min), aliquoted and either used directly to measure virus infectivity or stored at -80°C until use. To compare the infectivity of the Env variants, freshly prepared pseudoviruses were serially diluted and incubated with TZM-bl cells in the presence of 20 µg/ml DEAE-dextran. TZM-bl cells were supplied by John C. Kappes and Xiaoyun Wu (University of Alabama at Birmingham) through the NIH HIV Reagent Program. Forty-eight hours later, the luciferase activity in TZM-bl cells was measured using a luminometer.

Previously frozen pseudoviruses were used to evaluate the sensitivity of the Env variants to cold inactivation, antibody neutralization and inhibition by virus entry blockers. For these assays, similar TCID_50_ (50% tissue culture infectious dose) levels of the pseudoviruses were incubated with TZM-bl cells in the presence of 20 µg/ml DEAE-dextran. To evaluate the cold sensitivity of viruses with different Envs, pseudoviruses were incubated on ice for different lengths of time and virus infectivity was subsequently measured, as described above.

### Virus neutralization assay

Approximately 100-200 TCID_50_ of pseudovirus was incubated with serial dilutions of purified antibodies, sCD4-Ig, T20, BNM-III-170 or BMS-806 in triplicate wells of 96-well plates at 37°C for 1 hr. Then approximately 2×10^4^ TZM-bl cells in 20 µg/ml DEAE-dextran in medium were added to each well and the mixture was incubated at 37°C/5%CO_2_ for 48-72 hr, after which luciferase activity was measured. The concentrations of antibodies and other inhibitors that inhibit 50% of infection (the IC_50_ values) were determined by fitting the data in five-parameter dose-response curves using GraphPad Prism 8.

### Env expression and gp120-trimer association

HOS cells in 6-well plates were transfected with plasmids encoding His_6_-tagged Env variants and Tat at a ratio of 8:1, using the Effectene transfection reagent (Qiagen) according to the manufacturer’s instructions. Forty-eight hours after transfection, cells were lysed in buffer containing 1.5% Cymal-5. Clarified lysates were prepared, and a portion was saved as the “Input” sample. The remainder of the clarified lysate was incubated with Ni-NTA beads at room temperature or at 4°C for 1 hr. The beads were washed, boiled and analyzed by Western blotting with 1:2,000 goat anti-gp120 antibody (Invitrogen) and 1:3,000 HRP-conjugated rabbit anti-goat antibody (Invitrogen).

### Cholesterol depletion

To determine the IC_50_ of methyl-β-cyclodextrin (MBCD), approximately 100-200 TCID_50_ of pseudovirus was incubated with serial dilutions of MBCD at 37°C for 1 hr. TZM-bl cells were then added and luciferase activity was measured 48 hr later. The IC_50_ value of MBCD for each virus was then determined using GraphPad Prism 8, as described above. To determine the sensitivity of cholesterol-depleted viruses to antibodies and BNM-III-170, pseudoviruses were treated with MBCD at the IC_50_ for 1 hr at 37°C, after which the neutralization assay was carried out as described above.

### Activation of virus infection by sCD4-Ig and a CD4-mimetic compound, BNM-III-170

Pseudoviruses were incubated with CD4-negative, CCR5-expressing Cf2Th-CCR5 cells in 96-well plates. The plates were centrifuged at 1,800 rpm for 30 min at 21°C. Medium containing serial dilutions of sCD4-Ig or BNM-III-170 was then added. Forty-eight hr later, luciferase activity was measured.

## ACKNOWLEDGMENTS

We thank Ms. Elizabeth Carpelan for manuscript preparation. This work was supported by grants from the National Institutes of Health (AI145547, AI124982 and AI150471) and by a gift from the late William F. McCarty-Cooper.

## CONFLICT OF INTEREST

None.

